# The VR Billboard Paradigm: Using VR and Eye-tracking to Examine the Exposure-Reception-Retention Link in Realistic Communication Environments

**DOI:** 10.1101/2023.06.03.543559

**Authors:** Ralf Schmälzle, Sue Lim, Hee Jung Cho, Juncheng Wu, Gary Bente

## Abstract

Exposure is the cornerstone of media and message effects research. If a health, political, or commercial message is not noticed, no effects can ensue. Yet, existing research in communication, advertising, and related disciplines often fails to measure exposure and demonstrate the causal link between quantified exposure to outcomes because actual exposure (i.e., whether recipients were not only exposed to messages but also took notice of them) is difficult to capture. Here, we harness Virtual Reality (VR) technology integrated with eye tracking to overcome this challenge. While eye-tracking technology alone can capture whether people attend to messages in their communication environment, most eye-tracking research is bound by laboratory-based screen-reading paradigms that are not representative of the broader communication environments in which messages are encountered. Emerging eye-tracking field research suffers from an inability to control and experimentally manipulate key variables. Our solution is to measure eye-tracking within an immersive environment in VR that resembles a realistic message reception context. Specifically, we simulate driving down a highway alongside which billboards are placed and use VR-integrated eye-tracking to measure whether the drivers look at individual billboard messages. This allows us to rigorously quantify the nexus between exposure and reception, and to link our measures to subsequent memory, i.e., whether messages were remembered, forgotten, or not even encoded. We further demonstrate that manipulating drivers’ attention directly impacts gaze behavior and memory. We discuss the large potential of this paradigm to quantify exposure and message reception in realistic communication environments and the equally promising applications in new media contexts (e.g., the Metaverse).

## Introduction

Imagine driving down a seemingly never-ending highway, the billboard signs that line the road occasionally catching your attention. You briefly glimpse at some, examine others more closely, and completely bypass others. What will you remember when you reach your destination and why?

### Background

#### Exposure as the Cornerstone of Message Effects

Exposure is the cornerstone of media and message effects. As the concept suggests, exposure is about whether a message reaches a recipient, i.e., that the message enters a person’s information environment (1). This can include a sign along the road, a banner ad popping up while browsing the internet, or a commercial interrupting a TV program. It is obvious that if audiences do not receive a message, communication can not have any effect – just like a pill not swallowed cannot have any pharmacological effects.

Given the central role of exposure as a prerequisite of any message effect, much research has focused on measuring exposure. Virtually all media track information about audience sizes, such as newspaper readership, website visitors and browsing behaviors, TV and radio audiences, and so forth. Very often though, such data are only aggregate statistics, i.e., they provide information about average audience size but not whether individuals received a particular message. This is the difference between opportunities for exposure and actual exposure. Although conclusions based on exposure opportunities are possible (2), they are still subject to criticism.

Despite this key limitation, the evidence relating exposure to message effects is strong. For instance, opportunities for exposure (operationalized as how much a given message was “on air”) are directly related to the recognition of messages by target audiences (1,3,4). Similarly, several metrics of commercial messaging success, such as brand awareness or ad recall, are directly related to the volume of messaging, which is assumed to translate into exposure (5).

#### Encoded Exposure and Incidental Memory

To assess whether participants who might have been exposed to a message did actually receive it and encoded it into memory, researchers typically ask them to remember the content of the message (6). Different methods for probing memory exist, most notably free recall and recognition (7–9). In a free recall task, participants are asked to recall information they remember being exposed to. In contrast, the recognition method asks them to indicate whether or not an item was part of a set they encountered previously. Recognition is typically higher than recall because not all existing memory traces are retrieved during free recall.

Memory research has shown that multiple kinds of memory stores (e.g., explicit vs. implicit) exist and that different encoding operations affect performance (e.g., whether participants encode items superficially or more elaboratively) (10,11). The kind of memory most relevant to exposure research is incidental memory, which is memory formed without the intention to memorize. This is actually the default state we are in during much of our daily lives. Under such circumstances, attention is typically deployed to items that are intrinsically salient or relevant, and the resulting incidental memories are most relevant for message reception and effects research.

Although most of our memories are formed while we are engaged in everyday activities, most memory research focuses on more deliberative memory tasks and studies memory formation under laboratory conditions. This approach has led to important insights, but critics have long demanded that memory be studied under more ecological conditions and that more focus be placed on everyday memory (12–14). However, doing so requires overcoming obstacles that favor laboratory research and make naturalistic memory research challenging. These include that it is exceedingly difficult to study peoples’ behavior in natural environments and the difficulty of manipulating experimental factors therein. This applies to memory research in general, but also to research specifically focusing on memory for messages as it is studied in communication and advertising (6,15–18).

In sum, strong evidence underscores the importance of exposure for message effects. However, while aggregate-level exposure data are consistent with a dose-response relationship between exposure and recall, it is undoubtedly the level of a single exposure of an individual to a given message that causally underlies these effects. Said differently, the actual exposure occurs not just when a given message is “in the information environment”, but when it meets the eyes or ears of its recipients (1,19). However, not all messages we are actually exposed to are remembered, and we know shockingly little about how many messages we encounter in our daily lives. Therefore, there is a need to *close the measurement gap between opportunities for exposure, actual exposure, and memory*.

#### Eye-tracking for Measuring Exposure: Strengths and Current Limitations

Eye-tracking is an important tool for measuring visual information sampling (20). Eye tracking provides direct information about where an individual is looking, which is in turn related to what messages a person is paying attention to and how effectively they are processing the information. Due to these desirable characteristics, eye tracking is actually widely used to study how individuals respond to messages, where and for how they look, and so forth (21–24). However, one downside of previous eye-tracking studies is that they were confined to controlled laboratory environments and most eye-tracking was done using screen displays. While this approach is valid for studying how people browse internet websites, it does not allow for eye-tracking studies in more natural environments, such as highways, streets, and other contexts (25). As a result of this limitation, we know relatively much about situations in which people are placed in front of screens to study which displayed messages they attend to, but very little about more unconstrained information environments in which people freely initiate and terminate exposure to messages. However, to the extent that these situations are the natural norm rather than the exception, our knowledge and theories about exposure and its transmission into message effects are woefully incomplete.

Recently, wearable eye-tracking technologies have been developed that enable researchers to study message reception in natural contexts (25–27). While these technologies can overcome the limitations of screen-based eye tracking, they suffer the challenge of all field research, which is they don’t afford the experimenter to control the situation, offering limited potential for causal manipulation of variables that are assumed to influence outcomes.

#### VR’s Virtues: Realism, Control, and Measurement Capability

Virtual Reality (VR) can create a highly immersive and interactive experience, allowing researchers to accurately simulate real-world environments and study human behavior in a controlled and systematic manner. This potential of VR has been documented in various research contexts, including clinical psychological research, communication and advertising, as well as memory and navigation research, to name but a few (28–31). Key characteristics that recommend VR for research use include its realism, its opportunities for experimental control, and its potential to integrate measurement (32). We will next expand on each of these beneficial characteristics.

First, VR can create realistic environments that mimic real life – whether it is riding a rollercoaster, walking along a virtual plank, or driving down a highway. Visual information is particularly central to the human mind/brain and is one of the primary channels of human information intake that VR can virtually simulate. Thus, researchers can design virtual environments that mimic the essential appearance of a wide variety of human visual environments, and then explore the cognitive and socio-emotional mechanisms generated in visual environments by VR.

Second, because VR environments are virtually created, they can be precisely controlled. For instance, if the goal is to place a particular billboard along a virtual highway, researchers can create a virtual highway model and place a virtual billboard along the roadside – no permit or construction costs are required. This is obviously a great asset in terms of experimental control, which is widely seen as one of the most critical features to experimentally demonstrate the causality of theoretical variables (mechanism and intervention potential). Critically, however, in many situations, control is difficult and expensive to achieve (e.g., permit, cost), sometimes even completely impossible. In this sense, the ability to virtually create and manipulate experimental situations offers researchers a unique tool that optimally balances natural realism and experiment control.

Third, another desirable characteristic of VR is that it is relatively easy to integrate measurements for behavioral research: Because the virtual environment a user enters is fundamentally digital, a lot of data are naturally tracked as variables (e.g., a user’s position in the environment, head orientation, speed of movements). Additional variables can also be tracked (e.g., position of the hands). Currently, we see a strong trend to incorporate bio-behavioral metrics into VR systems (e.g., hand- and facial tracking for user interaction and avatar expressiveness, heart monitoring, eye-tracking, etc.) (33–35).

These three characteristics of VR – realism, control, and measurement potential – suggest it is an ideal candidate for research on the exposure-reception-retention link, especially if paired with eye-tracking. Indeed, some prior research has already used VR in this way. Many promising VR-related applications are proposed for related research purposes, although we are not aware of direct applications focusing on exposure to visual communication messages (33,36–40).

However, a fourth aspect should not go unnoticed. VR is heralded as the communication medium of the future, i.e., as an emerging media channel rather than just a methodologically advantageous gimmick (28,29,41,42). If true (see e.g., the rebranding of the social media company FaceBook to Meta), VR might be on the way to becoming a messaging environment in and of itself. In other words, if people are going to spend time in VR, they are exposed to a variety of messages while inside VR. Just like social media metrics (e.g., likes, comments, page impressions, and viewable impressions) have enabled the quantitative study of message diffusion on social networks, this development could also create opportunities to connect data about quantified individual-level exposure (e.g., fixation to a message) to subsequent outcomes.

### The Current Study and Hypotheses

The current study examines how quantified exposure to messages in a realistic environment relates to incidental memory of the messages. We introduce a novel VR billboard paradigm that simulates a drive down a realistic highway alongside which billboards are placed. Moreover, we combine this realistic and controllable VR environment with eye-tracking, thereby leveraging the integrated measurement potential of VR to capture exposure as it occurs.

To experimentally demonstrate the potential of this novel paradigm, we instructed half of the participants to look out for trash placed alongside the highway. The other half was instructed to look freely while driving down the highway. It is well documented that such a parallel, the attention-consuming task will distract participants and should lead to fewer fixations to the billboard messages.

Beyond checking how the competing task affects fixations to the billboards, we were primarily interested in whether looking at individual billboards would predict subsequent memory. The abovementioned reasoning on the exposure-retention link predicts that messages that were looked at should be committed to memory.

Finally, we wanted to explore general patterns of participants’ viewing behavior in this situation. To this end, we conducted additional data-driven analyses to identify patterns that would be predictive of outcomes.

## Methods

### Participants

Forty participants (*m_age_* = 25.6, *sd_age_*= 11.2; 18 female) were recruited from a study pool and via word of mouth. The local review board approved the study, all participants provided written informed consent to the protocol, and student participants received course credit. The sample size was set *a priori* to 40 participants, which was chosen based on power considerations and prior work in basic memory and VR research. Specifically, we determined that for an assumed large effect (*d* = 1.2), a sample size of 16 per group would be sufficient for high power (*1-*β = 0.95, α = 0.05) to detect a between-group difference in the number of recalled billboards. We rounded this number up to 20 per group. One additional participant whose goggles did not fit under the VR HMD was immediately replaced, resulting in a final sample of 40 participants.

**Fig. 1.**
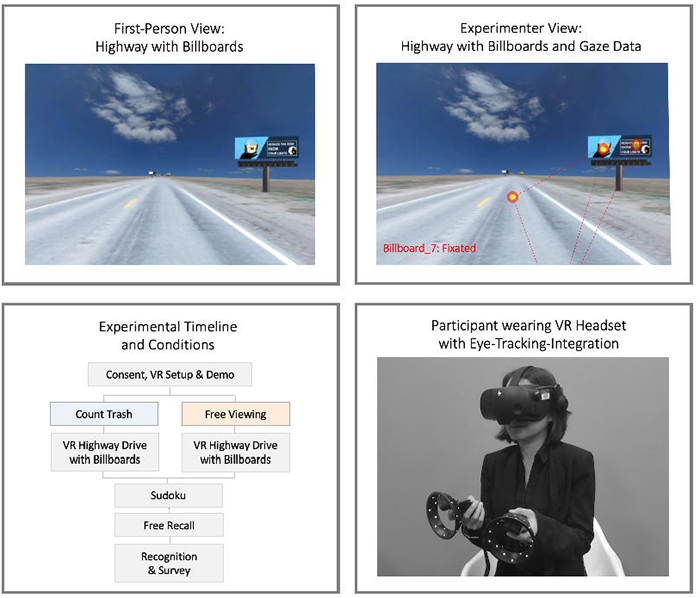
Overview of the experimental protocol. Top row: Screenshots of participant and experimenter views with one example billboard (‘drunk_driving’). Bottom left: Experimental timeline and conditions. Half of the sample was instructed to count trash along the road, the other half was instructed to simply drive down the highway. Except for this difference in instruction, the highway and billboards were identical for both conditions. Bottom right: Participant wearing an HP Reverb G2 Omnicept VR headset.

### Materials and Equipment

#### VR highway environment and billboard signs

We developed a virtual highway in which 20 billboards and additional highway-typical elements (construction signs, empty soda cans) were placed along the road. The core highway model was downloaded from Sketchfab.com and consisted of a digitized 3D model by the Nevada DOT (43). It featured a straight stretch of highway 50 taken near Cold Springs. Virtual billboard signs were placed along the road using 3D-billboard model stands, and the billboard messages were assigned to each of the 20 billboards stands in a programmatic fashion (randomized across participants). The distance between successive billboards was assigned randomly and then kept fixed across participants.

#### Billboard messages

We developed 20 visual billboard messages using templates from Canva.com. Half of the billboards were about health-related topics (e.g., drinking, vaping, smoking, marijuana, seatbelt use, and distracted driving). The second half of the billboards were typical advertisements (e.g., retail, lawyer services, hotels, and restaurants/food). The billboard messages all featured basic imagery along with some text, and their design was deliberately kept relatively simple but still typical of the kinds of billboards present on US highways (see Fig. 1 and Supplementary Materials).

#### VR and eye-tracking

We relied on the Vizard VR software to create the VR environment, run the study, and track user behavior, including eye-tracking measurements (Vizard, 7.0; (44). The VR device was an HP Reverb G2 Omnicept that includes eye-tracking capabilities. Participants used the right VR controller to accelerate and drive forward along the virtual highway. Because the highway was perfectly straight, no steering was required.

### Experimental Procedure and Conditions

Once participants arrived and consented to the study, they completed a quick vision test, put on the VR headset, and underwent a calibration routine. Next, the participant entered a demo version of the study to familiarize them with VR, the virtual space, and the navigation. Then, the main experimental session was started, which involved driving down the virtual highway. Half of the participants were instructed to count the number of trash items in the environment (distraction condition). The other half were told to explore the environment while driving down the highway freely (free-viewing condition).

After completing their virtual drive (which took about 10 minutes), participants were given a set of Sudoku puzzles for 2 minutes. Then, the experimenter conducted a structured interview that asked participants about the number of trash items they saw, their general virtual driving experience, and which billboards they recalled (free recall task). As the last step of the study, participants completed an online questionnaire via Qualtrics that collected demographics, their experiences with the VR technology, and recognition of billboards. Specifically, we asked participants to report on their experience of spatial presence and the occurrence of symptoms while in VR (45); (46). For the recognition test, they were shown the 20 experimental messages and four distractors and asked whether they remembered seeing the messages during the highway drive. The purpose of the distractors was to gauge participants’ tendency to recognize all messages as seen. Finally, participants were debriefed, and their eye-tracking data was saved.

#### Main Measures and Analysis Methods

The main variable measured during the virtual drive was participants’ fixation on a given billboard, which was detected algorithmically and saved to disk in a spreadsheet. The fixation threshold was set to 0.25s. Thus, for every participant, the virtual drive yielded a spreadsheet containing where a given banner was fixated (and how often, e.g., time 15s, fixation, billboard_1, etc.).

Because billboard images were randomly assigned to individual billboard sign positions, a python program was written to resort the individual images to a given participant’s eye-tracking data (e.g., time 15s, billboard_1, drunk_driving.jpg, …), allowing for subsequent data aggregation across participants and billboard messages.

The recall data (information on whether a participant brought up the billboard during the free recall task, e.g., “I recall seeing a billboard about drunk driving”) was merged with the fixation information, and so was the recognition data (information on whether a participant recognized the billboard at the end of the study from a list of billboard images).

We document the analysis and provide code in the study’s online repository [link to data and scripts on OSF and Github will be included in the final manuscript]

## Results

We measured eye gaze while participants drove down a virtual highway with billboards alongside. One group was instructed to pay close attention to trash, another group could explore the environment while driving. Once they reached the end of the highway, we tested the participants’ incidental memory for the billboard messages (they had not been told that they would be asked about the messages).

Participants’ verbal comments about the study, collected during the verbal interview, revealed that they found the virtual highway drive realistic and engaging. The post-experimental survey data confirms these observations. Specifically, participants reported experiencing spatial presence in the VR environment (*mean_spatial presence_*= 3.8; range 1-5, i.e., all items above the scale midpoint). Participants also reported almost no symptoms (e.g., dizziness, fatigue, or eyestrain; *mean_VR-symptoms_*= 1.36, range 1-4, i.e., all items below the scale midpoint).

### Fixations as a Function of Load Condition and General Memory Performance

To examine the effect of the experimental conditions (trash-counting vs. free-viewing), we compared the number of fixations to billboards. As predicted, we find that participants in the free-viewing condition had significantly more fixations on the billboards than participants in the trash-counting condition where participants’ attention was directed more to the road than the billboards (*mean_fixations:free-viewing_* = 52.8, *sd* = 22.9; *mean_fixations:trash-counting_* = 21.8^1^, *sd* = 16.1; *t_38_* = 4.98, *p* < 0.001; *d* = 1.58). These results are shown graphically in Fig. 2 (left panel).

Next, we examined the memory performance collected in the interview. In the free recall test, participants in the free-viewing condition recalled an average of 6.45 billboards (*sd* = 1.57, *recall_rate_free-viewing_* = 0.32, significantly more than the average 2.95 (*sd* = 1.76, *recall_rate_trash-counting_*= 0.15) billboards the participants in the trash-counting condition recalled (*t_38_* = 6.63; *p* < 0.001; *d* = 2.1; see Fig. 2, middle panel).

Carrying out the same analysis on recognition data revealed even more pronounced results: Participants in the free-viewing condition recognized on average 14.6 billboards (*sd* = 3.25; *recognition_rate_free-viewing_* = 0.73) compared to only 7.1 billboards recognized in the trash-counting condition (*sd* = 3.68; *recognition_rate_trash-counting_* = 0.35), which is a highly significant difference (*t_38_* = 6.78, *p* < 0.001, *d* = 2.14, see Fig. 2, right panel).

**Fig. 2.**
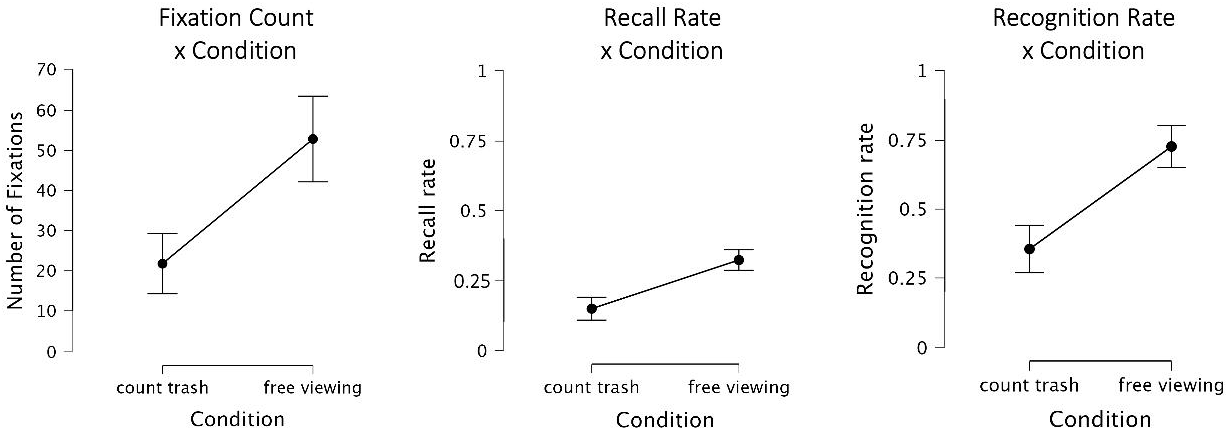
Number of fixations, free recall, and recognition rates by condition.

### Relationship between Fixations (Exposure and Attention) and Memory

Next, we focused on the relationship between fixations on individual billboards and subsequent memory for the billboards. Toward this end, we determined for every participant the number of looked-at billboards (i.e., fixated at least once) that were later recalled (or recognized) and the corresponding number of billboards that were not-looked-at (i.e., never fixated). The resulting table was then submitted to an ANOVA, which revealed highly significant and consistent effects for both ways of assessing memory.

For free recall, there was a highly significant interaction effect between condition and fixation status (*F* (_1,38_) = 25, *p* < 0.001, η*^2^_p_* = 0.4) and a highly significant main effect of fixation status (*F*(_1,38_) = 132.6, *p* < 0.001, η*^2^_p_* = 0.78). Follow-up tests confirmed higher recall in the free-viewing condition.

The results for the recognition data closely resembled the recall analysis: A highly significant main effect of fixation status (*F*(_1,38_) = 91.2, *p* < 0.001, η*^2^_p_* = 0.71) was qualified by a significant ordinal interaction of condition and fixation status (*F_(1,38)_*= 23.5, *p* < 0.001, η*^2^* = 0.38). Follow-up tests again confirmed that recognition memory was higher in the free-viewing condition compared to the trash-counting condition. In other words, we find that if a billboard is looked at at least once, this boosts the likelihood it will be remembered by a factor of 5-20 (depending on the condition and how memory is measured).

**Fig. 3.**
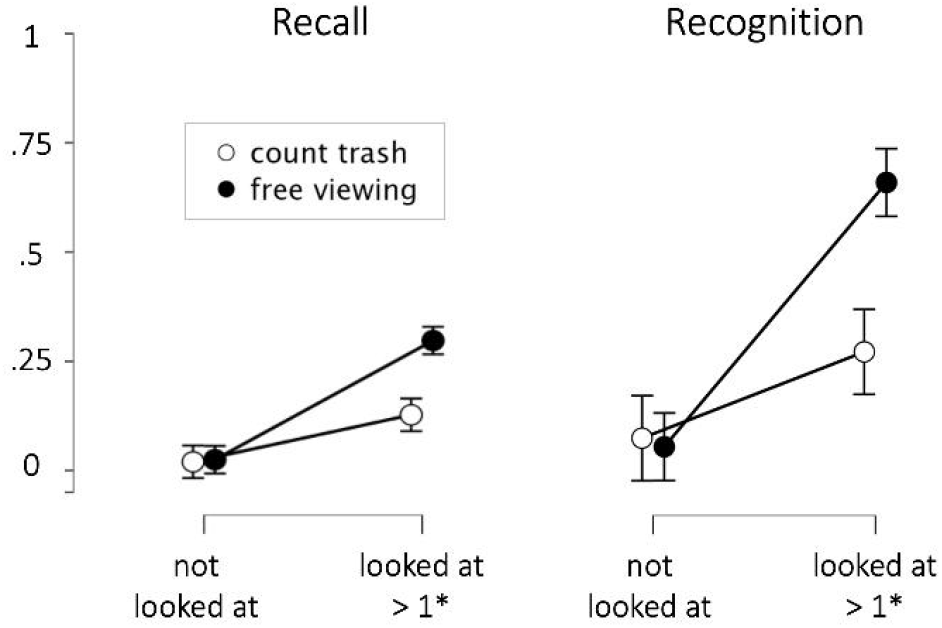
Relationship between fixations and subsequent message memory. Probability of recall (left) and recognition (right), based on whether a billboard was looked at (not fixated or fixated at least once) and condition (count trash vs. free viewing).

To illustrate this more clearly, we created a Fig. that jointly visualizes whether a billboard was looked at and whether it was recalled or recognized, respectively, and in which condition (see Figs. 3 & 4). As can be seen, in the trash-counting condition (blue dots), many billboards are not looked at at all. In the free-viewing condition (orange dots), more billboards are looked at (see results in the previous paragraph). Critically, however, the billboards that are never looked at are practically never recalled (top left quadrant in the scatter plots, see discussion for explanation). The banners that were looked at (right column), are far more often recalled, and almost always recognized.

**Fig. 4.**
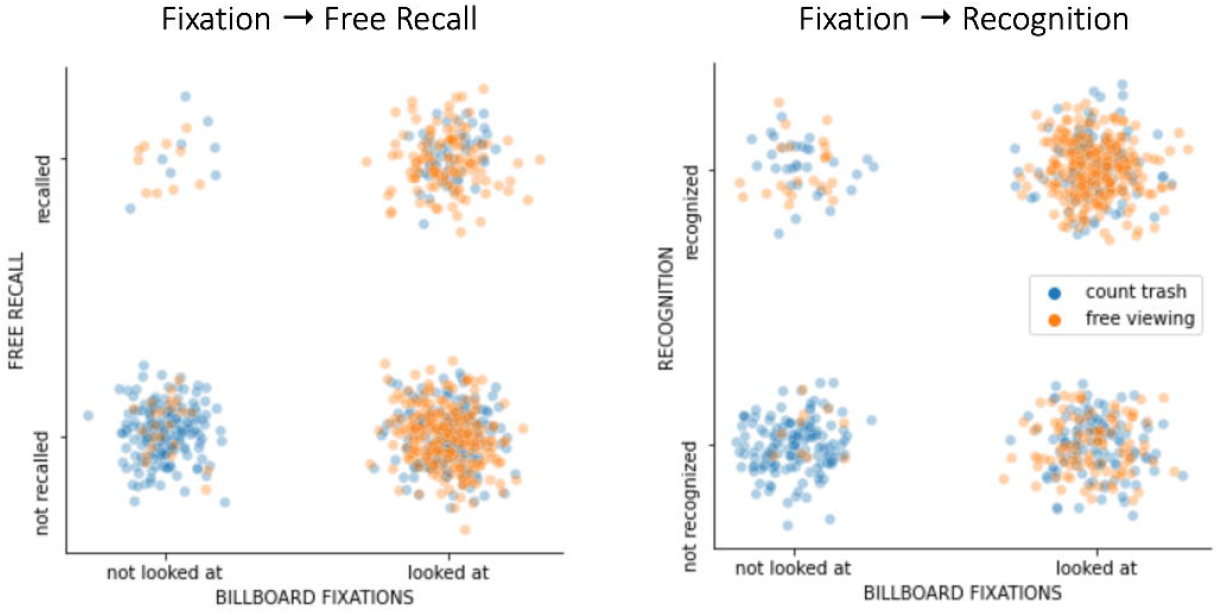
Relationship between fixations and subsequent message memory at the level of single messages. Left panel: Fixations and free recall performance. Every dot represents one billboard, color-coded based on whether participants were instructed to count trash (distraction) or view freely. Note that dichotomous variables (0 – not looked at/not recalled, 1-looked at/recalled) were jittered randomly to aid visualization. Right panel: Same analysis but based on a recognition memory test.

To examine this strong contingency between looking and remembering at a more fine-grained level, we further unpacked the fixation data. Specifically, we extracted for every participant whether a billboard was never looked at, looked at a few times (i.e., at least once but less than that participant’s medium fixation count across all 20 billboards), or looked at often (more than that participant’s medium fixation count across all 20 billboards). The results of this analysis are illustrated in Fig. 5, and they are statistically significant. A repeated measures ANOVA for the average number of items recalled (DV) revealed a strong effect of Viewing Behavior Intensity (*F_(2,76)_* = 12.1, *p* < 0.001, η*^2^* = 0.24) with a significant interaction of Viewing Intensity * Condition (*F_(1,38)_* = 3.68, *p* < 0.05, η*^2^* = 0.09).

**Fig. 5.**
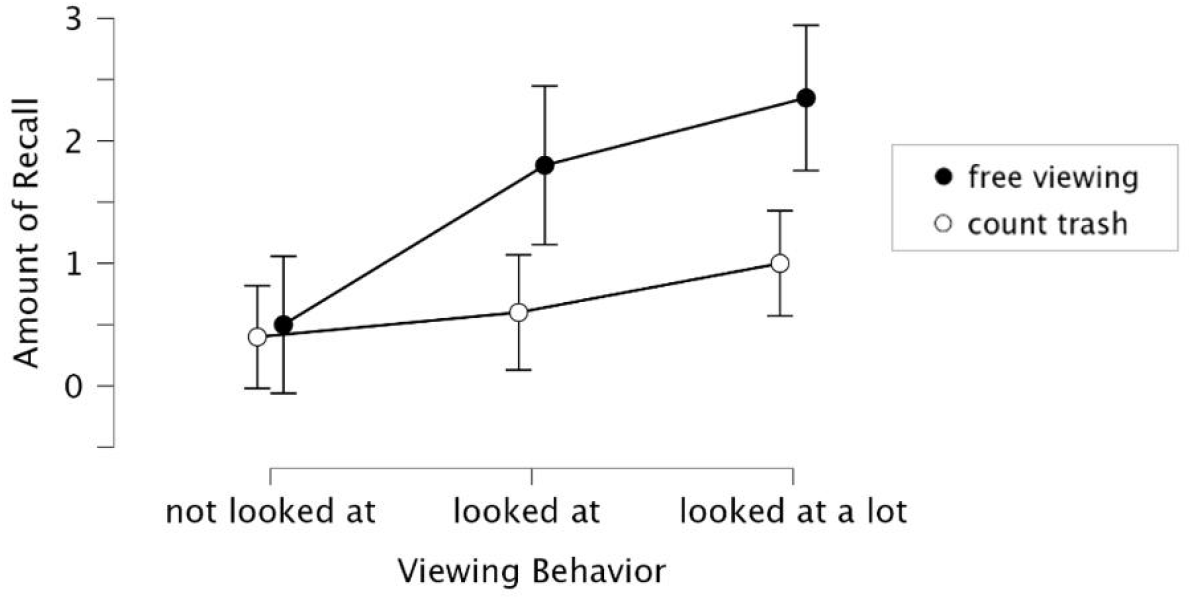
Relationship between Viewing Behavior Intensity and Message Recall at a more refined level (i.e., beyond looking vs. no-looking).

#### Exploratory Analyses

The approach presented here affords predictive modeling. To this end, we used scikit-learn (47) to create a model that could classify whether a billboard would be recalled or not based on the existing variables, i.e., which billboard was presented (e.g., buckle-up, drunk_driving, hotel, etc.), how often the participant fixated it, in which position the item was viewed, and the condition (trash-counting vs. free-viewing). Using a 5-fold cross-validated SVC prediction, we found that this simple model performed well, with a ROC-AUC score of 72.8% – compared to 50% for a dummy classifier (note that we used penalization to deal with the imbalance classes, i.e., recall being rarer than no recall). Said differently, once we know that a participant looked at a given billboard, we can predict more accurately whether this participant will later recall it. This relationship can also be derived from the statistically significant effects and the data shown in Figures 3-5.

In addition to statistical testing and predictive models, we carried out additional analyses to examine false recognition, results for individual participants and individual billboards, effects of item position, and health vs. commercial billboard content.

To gauge the degree to which participants would be prone to false recognition, we included distractor billboards in the recognition set (i.e., billboards that were never seen). However, these distractors were only rarely falsely recognized, significantly less than all presented billboards, and only one participant mis-recognized more than two distractor billboards. Thus, even though recognition measures can be prone to guessing, this does not seem to be the case here.

We also explored the relationship between fixations and memory and between different memory measures at the individual level: In the trash-counting condition, the number of fixations and memory measures were highly correlated (*r values* > 0.7, *p values* < 0.001), suggesting that participants who were more interested in the billboards or the study also remembered them better. In the free-viewing condition, this was not the case (*r values* were nominally even negative). In both conditions, however, recall and recognition were positively correlated (*r* = .54, *p* < 0.001 for the trash-counting condition, *r* = 0.2, *n.s.,* for the free-viewing condition). While these results are interesting and point to effects of motivation or interest, we opted not to investigate them further because the current sample was relatively small for studying individual differences.

Moreover, we inspected the potential influence of the billboards’ position (beginning vs. middle vs. end) on the probability of fixation, recall, or recognition. However, we did not find such effects, nor evidence of an interaction with the condition. In both conditions, position curves were parallel and flat.

Inspection of the results for individual billboards, however, revealed interesting effects: Specifically, as shown in Fig. 6, some items were often recalled (e.g., buckle_up, disobey_vape, and burger) – others were barely remembered. This is also consistent with the predictive modeling result, where adding the item (one-hot-encoded) as a feature increased accuracy. Most likely, this is due to intrinsic differences between the billboards – either because of the topic’s relevance to participants or because of low-level physical differences, such as saliency. Of note, we did quantify perceptual saliency (48) but did not see a significant relationship with memorability.

Lastly, we also compared the health-related banners against the commercial banners, finding no significant differences. Nominally, health-related billboards were slightly more often recalled (also see Fig. 6), but the effect was insignificant (*F_(1,36)_* = 3.53, *p* = 0.07). Across both conditions (independent participants), the same billboards tended to be recalled more often, as indicated by a significant vector correlation between trash-counting and free-viewing (*r* = 0.82, *p* < 0.001).

**Fig. 6.**
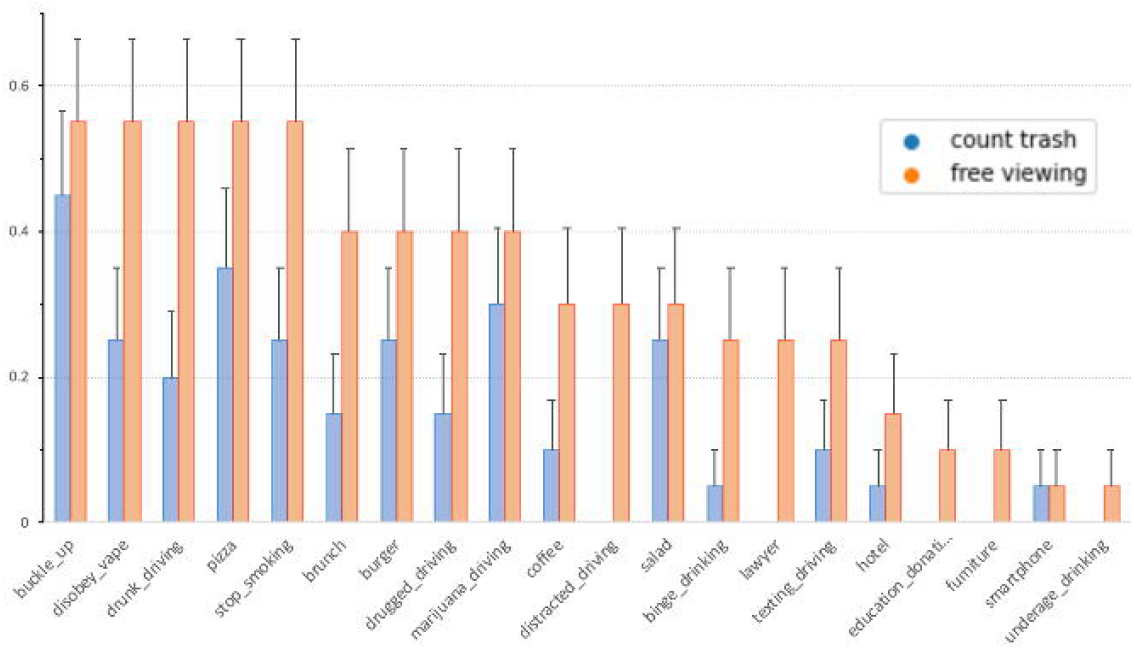
Analysis for individual billboards. Across both conditions (independent participants), the same billboards tended to be recalled more often.

## Discussion

Messages are intended to inform and influence recipients. However, this requires that they are viewed, i.e., that audience members are actually exposed to the message. Therefore, exposure is the cornerstone of all message effects, but measuring exposure is challenging – especially at the individual level and within realistic messaging contexts. Here we created a VR paradigm that immerses users in a realistic environment familiar to many: a drive down a highway with billboard signs along the road. Using a VR-integrated eye-tracker, we recorded whether participants looked at individual billboards and we link this information to subsequent memory for the billboards. Our results show that this approach allows us to rigorously assess the exposure-reception-retention nexus.

### Discussion of Main Results

The current results are very clear and straightforward: The VR Billboard Paradigm enables studying whether people look at (i.e., take in the information) from the messages they were exposed to. As simple as this sounds, the significance of it becomes apparent if one considers that exposure is the cornerstone of message effects, but exposure is often only inferred rather than actually measured (i.e., how often a TV ad is on air and typical audience sizes are taken as opportunities for exposure). Clearly, these indirect ways of assessing exposure miss the point because what really matters for message effects is *actual reception*, not *fiat exposure* (“Let’s hope people will look at the message”). Our paradigm now makes it possible to study this and to do so in a way that strikes a balance between realism and experimental control.

Perhaps the most important effect is that participants’ viewing behavior was significantly associated with message memory: Technically, one could have argued that all participants passed by all messages (i.e., had the opportunity for exposure). However, measuring their visual information sampling via eye tracking made it possible to measure actual exposure – and thus reception -, and this explained whether billboards would be recalled.

A third point was the strong influence of the task: Participants who were instructed to drive freely looked at the billboards more often and they recalled them more often. By contrast, participants who were instructed to count trash along the road showed very few fixations towards the billboards and generally low memory. Again, this perfectly matches our predictions that participants’ attention would be consumed by the task, as it is well-known that attention and memory are tightly coupled (49–52).

These results all support our main argument, which is that exposure and reception are the prerequisites of message retention (memory). The present approach thus has value for pinpointing the mechanism that leads from exposure to retention: Specifically, the causal chain starts with the presence of a message in the information environment (opportunity for exposure), then the person noticing and taking in the message (actual exposure, reception), to subsequent memory (retention). Further evidence for this causal pathway is also provided by the dose-response relationship, i.e., the marked differences between fixations and memory in the trash-counting vs. free-viewing conditions, and by the fact that messages that were looked at more were recalled more often.

### Broader Implications

The new approach presented here holds significant value for understanding exposure and reception as the critical nexus between message and receiver in communication. As such, the approach is not only methodologically intriguing but also promises to advance our understanding of the theoretical factors that affect the exposure-reception-retention nexus.

Although in communication science the concept of exposure has remained hard to study naturalistically, experimental memory research is an area in which exposure has always been manipulated – by forcing participants to attend to messages and then study the effects on memory. As such, laboratory work on memory encoding and work flowing from incidental and ecological memory perspectives is complementary to the current approach (52,53), although our emphasis differs by taking a communication perspective (3). For instance, in memory research, the levels of processing framework highlighted how recall of memoranda varies based on processing depth (10,54). The core assumption is that the mental operations carried out over items (e.g., whether they are processed semantically or only superficially) influence the probability of recall. Such work has also found its way into communication science, for instance via the popular elaboration-likelihood-model and related work (55,56). Likewise, the concept of involvement in advertising has been proposed to refer to the degree of personal connections message recipients make with a message once they received it (57). Finally, the notion of exposure states also points to the importance of examining the psychological processes message recipients engage in once they are exposed to messages (19). Thus, these different models and theories all have in common that they require measuring i) whether messages are received and ii) how people engage with them. The VR billboard paradigm presented here can definitely ascertain the former (whether messages are seen). To the extent that fixation amount and length can give insight into the latter (how messages are engaged with), we can also examine this with the current paradigm. Moreover, the paradigm can easily be expanded to measures like pupil dilation (or derivative metrics like fixation length, paths, etc.). In sum, the VR billboard paradigm resolves a longstanding problem in a new way that promotes method-theory synergy (58) between VR and eye-tracking research, laboratory, and everyday memory, and work on the exposure-reception-retention nexus in communication.

Beyond these theoretical considerations, this approach clearly has significant applied potential as well: First, the VR billboard paradigm is directly applicable to billboard advertising in the real world. For instance, it could be immediately used to empirically examine the effects of new constructions on existing billboards (e.g., as legal testimony), forecast billboard effectiveness, and so forth. Second, the approach can easily be adapted to other applied messaging questions because many message delivery contexts could be implemented in an equivalent manner. These include all forms of outdoor advertising, including airports, public transportation, and public spaces like Times Square in New York, the strip in Las Vegas, or any place where large audiences pass by. In all of these cases, the ability to experimentally manipulate key characteristics of the appearance or the users’ state and assess the effects of such manipulations on quantified user behavior (here, eye-tracking) could be of major value.

We note that we are not the first to point to the potential of VR and eye-tracking for studying exposure and memory and that several related works exist. For instance, Kim et al. (59) have suggested a 360-degree video paradigm for measuring viewing behavior in naturalistic settings (i.e., 360-degree videos of real cityscapes). This approach combines realism and eye-tracking. From an experimental point of view, however, the ability to control the placement and content of billboards, or even make message delivery contingent on behavior, offers key strengths and innovations.

Going forward, we also expect key advances by integrating additional measures beyond the current eye-tracking. For instance, our results here focus on the eye gaze fixations and make hardly any use of pupil dilation or heart rate, both of which are already integrated into the HP Reverb G2 Omnicept headset. Kim et al. (59), for instance, did combine their video with MRI measurement. Although VR is challenging to combine with MRI (because the equipment is not compatible with brain scanners and head motion presents problems for MRI), other options exist and will likely become more widespread. These include EEG and fNIRS, which can provide additional insights into, e.g., the neural basis of memory formation and attention (51,60,61). We also note that there were very few messages that were not looked at, but were still remembered (very rarely recalled freely, but sometimes recognized, see Fig. 4). This can be explained by parafoveal or ultra-fast vision (i.e., below fixation threshold, 62-64) and one could argue that these events are rare. Still, in such a case, neural measurements could add in information beyond eye tracking alone.

### Strengths, Limitations, and Avenues for Future Research

Key strengths of the VR billboard paradigm include that it is simple, realistic, flexible, and scalable. Using VR in combination with eye-tracking to study message reception is not confined to billboards on highways, however, but could be applied to other settings. It would, for instance, be very simple to exchange the environment and use the same available python code to detect fixations on messages in e.g. city settings.

Perhaps the biggest advantage of this approach over existing work (either screen-based eye-tracking or eye-tracking field research) is that it allows for the controlled testing of causal mechanisms, while preserving a high degree of realism. The ability to measure precisely and objectively and to control variables experimentally are the key prerequisites for causal mechanism identification in, e.g., the biological and behavioral sciences. In the social sciences, which often rely on macro-level association data, these features are difficult to achieve. In this sense, the current paradigm holds great potential to overcome many limitations that have plagued message exposure research. Of note, though not the main focus of our study, this paradigm would seem equally promising for applied memory research (52,53).

Like all research, the current study has several limitations. One limitation is that although the VR experience featured a realistic version of a real highway drive (a digital twin of highway 50 near Cold Springs), some elements of real life were missing (e.g., opposing traffic, birds, curves, and passages through towns, etc.). Likewise, our experimental messages are also limited in variety, number, or design- and content elements. We deliberately made these choices to balance experimental control and realism, but it could of course be argued that specific features might matter. Fortunately, it is easy to add and test such factors, and high-realism driving games demonstrate that this is feasible (e.g., the popular Need for Speed or GTA series).

Another limitation concerns the mostly student sample and its size. While our sample was adequate for the study’s goal, which was to demonstrate the value of this new paradigm by eliciting a fairly basic memory effect, future studies examining smaller or more contextual effects will require larger and more diverse samples. Given that most VR research is still conducted in laboratory settings and measuring one person at a time, this will lead to a bottleneck at the data acquisition stage. However, as VR enters the mass market, we can expect that VR crowd studies will emerge. This would then provide researchers with access to samples the size we see in survey research, but with the added opportunity to capture biobehavioral data during message reception.

Considering the above-mentioned strengths and limitations, we regard the following avenues for future research as promising: First, it would be promising to extend the context from billboards along highways to broader messaging contexts, like cityscapes, airports, and so on. All of these situations can be created virtually with little effort, and several free 3D models do exist. Similarly, even the current VR billboard paradigm offers a host of options. For instance, it would be promising to examine the influence of distractions or contexts, such as concurrent radio messages along the drive, or manipulations of user-state variables (like having hungry participants view food billboards; (65)).

Along these lines, we also see much potential for more dynamic manipulations. By this, we mean that the current study only manipulated static billboards and the messages that were shown along a virtual drive. The next step would be introducing manipulations in which the messages are contingently administered. For instance, it would be possible to show a message if the driver previously looked at another one or to show a message for as long as needed until the driver viewed it. These options show the enormous potential for persuasion and nudging strategies, which are of course a double-edged sword: On the one hand, these could be leveraged to improve the effectiveness of health communication. On the other hand, they could be used for commercial advertising. Regardless of the intent of the messenger, however, it is undoubtedly the case that such applications would bring communicators closer to the long-standing goal of being able to “give the right message to the right recipient, at the right moment in time.”

### Summary and Conclusion

In sum, we suggest a new paradigm to study the link between attention and retention, or exposure and memory for messages. The VR billboard eye-tracking paradigm allows for studying incidental memory formation under highly realistic conditions, but with exquisite experimental control and integrated bio-behavioral measurements. The result that fixations are related to memory confirms the link between exposure/attention and retention/memory, underscoring the potential for this paradigm to study memory in real-world contexts and communication effects in the new information ecosystem.

## Footnotes

1 Based on this average it is tempting to assume that all participants in the trash-count condition may have looked about once at every billboard (20 in total). However, this was not the case. Rather, a few participants looked are some billboards more often, and many participants in the trash-counting condition did not look explicitly at many billboards.

